# Naked mole-rats (*Heterocephalus glaber*) do not specialise on cooperative tasks

**DOI:** 10.1101/2021.03.22.436002

**Authors:** Susanne Siegmann, Romana Feitsch, Daniel W. Hart, Nigel C. Bennett, Dustin J. Penn, Markus Zöttl

**Affiliations:** Konrad Lorenz Institute of Ethology, University of Veterinary Medicine, Vienna, Austria; Mammal Research Institute, Department of Zoology and Entomology, University of Pretoria, Pretoria, Gauteng, South Africa; Ecology and Evolution in Microbial Model Systems, EEMiS, Department of Biology and Environmental Science, Linnaeus University, SE-391 82 Kalmar, Sweden

**Keywords:** Behavioural specialisation, division of labour, eusociality, cooperative breeding, helping, social evolution

## Abstract

It has been proposed that naked mole-rat (*Heterocephalus glaber*) societies resemble those of eusocial insects by showing a division of labour among non-breeding individuals. Earlier studies suggested that non-breeders belong to distinct castes that specialise permanently or temporarily on specific cooperative tasks. In contrast, recent research on naked mole-rats has shown that behavioural phenotypes are continuously distributed across non-breeders and that mole-rats exhibit considerable behavioural plasticity suggesting that individuals may not specialise permanently on work tasks. However, it is currently unclear whether individuals specialise temporarily and whether there is a sex bias in cooperative behaviour among non-breeders. Here we show that non-breeding individuals vary in overall cooperative investment, but do not specialise on specific work tasks. Within individuals, investment into specific cooperative tasks such as nest building, food carrying and burrowing are positively correlated, and there is no evidence that individuals show trade-offs between these cooperative behaviours. Non-breeding males and females do not differ in their investment in cooperative behaviours and show broadly similar age and body mass related differences in cooperative behaviours. Our results suggest that non-breeding naked mole-rats vary in their overall contribution to cooperative behaviours and that some of this variation may be explained by differences in age and body mass. Our data provide no evidence for temporary specialisation, as found among some eusocial insects, and suggests that the behavioural organisation of naked mole-rats resembles that of other cooperatively breeding vertebrates more than that of eusocial insect species.

## Introduction

Task specialisation among members of social groups is considered a hallmark of social evolution and can lead to improvements in group efficiency (Chittka and Muller 2009, Bourke 2011). The most extreme cases of task specialisation are found among social insects, where individuals show divergent developmental trajectories that lead to functionally different and morphologically specialised castes of workers (Wilson 1971, Bourke 2011). Other social insects show temporary specialisation in the absence of morphological specialisation, and workers pass through successive developmental stages that are characterised by temporary specialisation in specific tasks (Seeley 1982, Biedermann and Taborsky 2011, Mersch et al. 2013). In contrast to insects, group living vertebrates rarely show evidence of specialisation, and usually, individuals vary in their overall investment in cooperative tasks depending on the individual’s characteristics and environmental conditions (Cockburn 1998, Clutton-Brock et al. 2003). However, the social mole-rats of the family *Bathyergidae* may represent an exception among group-living vertebrates, and it has been controversially debated to what extent their social organisation resembles that of social insects groups (Jarvis 1981, Bennett 1990, Burda 1990, Crespi and Yanega 1995, Bennett and Faulkes 2000, Burda et al. 2000, Scantlebury et al. 2006, Boomsma 2013, Boomsma and Gawne 2018).

Early research on naked mole-rats (*Heterocephalus glaber*) has suggested that some non-productive individuals specialise permanently on specific work-related tasks and that variation in their cooperative behaviour is a consequence of the development of distinct castes – similar to those found in eusocial insects (Jarvis 1981, Jarvis et al. 1991). Variation in growth, body mass and behaviour were thought to be consequences of divergent developmental trajectories, where small-bodied workers specialise in acquiring indirect fitness benefits generated by helping related individuals, and large individuals were thought to maximise chances of direct reproduction by dispersing or replacing the breeder (Jarvis et al. 1991, O’Riain et al. 1996). Other studies suggested that variation in cooperative behaviour of naked mole-rats may instead represent temporary specialisation, similar to age-related polyethisms found in some social insects and that contrasts in behaviour may be explained by age-related changes of behaviour, where individuals pass through stages of development and conduct different tasks depending on their age (Lacey and Sherman 1991, Lacey and Sherman 1997, Faulkes and Bennett 2016). More recent studies have suggested that naked mole-rats show behavioural flexibility and that cooperative behaviour may be adjusted to the group composition and other social and environmental factors (Mooney et al. 2015, Gilbert et al. 2020). However, it remains unclear whether non-breeding naked mole-rats specialise temporarily on specific work-related behaviours, or whether individuals vary mostly in their overall commitment to cooperative behaviours (Thorley et al. 2018, Braude et al. 2021).

Evidence from other social mole-rat species challenges the hypothesis that specialisation is common in mole-rats. Several species of the genus *Fukomys* and *Cryptomys* show a similar social organisation to that of naked mole-rats and exhibit high reproductive skew and cooperative foraging, though their groups are usually smaller than naked mole-rat groups (Bennett and Jarvis 1988, Bennett 1990, Jarvis and Bennett 1993, Jarvis et al. 1994). Whereas subordinate Damaraland mole-rats (*Fukomys damarensis*) exhibit differences in their overall investment in cooperation and show age and size-related changes, the individuals do not specialise in specific tasks, and behavioural variation appears to be a consequence of differences in age, growth and body condition among non-breeders (Bennett and Jarvis 1988, Bennett 1990, Zöttl et al. 2016a, Thorley et al. 2018, Torrents-Ticó et al. 2018a). Similarly, research on the cooperatively breeding Micklem’s mole-rat (*Fukomys micklemi*) showed that non-breeding individuals lacked task specialisation (Van Daele et al. 2019) and radio-tracking studies of free-living Ansell’s mole-rats (*Fukomys anselli*) did not find evidence for behavioural specialisation (Šklíba et al. 2016). However, sociality has evolved independently in naked mole-rats and their relatives of the genera *Fukomys* and *Cryptomys*, and it is possible that patterns of behavioural organisation differ as a result of larger mean group sizes in naked mole-rats (Bennett and Faulkes 2000, Faulkes and Bennett 2007, Visser et al. 2019).

To demonstrate behavioural specialisation, it is necessary to show that individuals trade-off investment in different forms of cooperative behaviours. This would be expected to generate negative correlations between some cooperative behaviours across individuals over a considerable amount of time (English et al. 2015, Thorley et al. 2018). Previous studies of naked mole-rats have sometimes suggested that specialisation occurs on the grounds that individuals of different body mass or age show contrasts in their investment in specific tasks (Jarvis 1981, Jarvis et al. 1991, Lacey and Sherman 1991, Lacey and Sherman 1997), or by showing that different forms of cooperative behaviours load on different axes in principal component analyses (Mooney et al. 2015). However, these patterns do not necessarily imply specialisation on an individual level, and it remains unclear to what extent naked mole-rats specialise in cooperative tasks.

In this study, we investigated whether non-breeding individuals in captive naked mole-rat groups specialise across three different cooperative tasks, which are burrowing related activities, nest building and food carrying. To do this, we collected longitudinal behavioural records of 169 marked individuals in 11 groups using an instantaneous sampling protocol and analysed the behavioural frequencies with multilevel, multinomial logistic regressions. These generalised linear mixed models are logistic regressions that allow the estimation of within-individual correlation, while also estimating the effects of individual characteristics and environmental effects on behavioural variation (Koster and McElreath 2017, Thorley et al. 2018). Trade-offs between different cooperative behaviours at the individual level would result in negative individual random effects correlations. In contrast, positive correlations would indicate that individuals that perform one cooperative task are also much more likely to perform another kind of cooperative task more frequently.

We also investigated whether the expression of cooperative behaviour of naked mole-rats is predicted by individual characteristics (body mass, age) and group size, and whether there is a sex bias in the expression of cooperative behaviour. Variation at these levels and behavioural specialisation are non-mutually exclusive phenomena and do not preclude each other. As such, when addressing questions about behavioural specialisation it is important to include the effects of individual characteristics and group level traits because divergent behavioural trajectories in different tasks may reflect the relative costs and benefits of specific cooperative behaviours at different developmental or life-history stages (McNamara and Houston 1996, Taborsky and Grantner 1998, Heinsohn and Legge 1999, Clutton-Brock et al. 2003).

## METHODS

### Animals and housing

The study includes data from five groups of naked mole-rats housed at the Vienna Zoo (Tiergarten Schönbrunn) in Austria, and six groups housed at the University of Pretoria in South Africa, with group sizes ranging from 12 to 45 individuals. All animals were born and raised in captivity and housed in tunnel systems made of either transparent PVC or glass. Each group occupied a self-contained tunnel system (3.20–7 m) including at least one nest box and one toilet area. Temperatures in the housing facilities were maintained close to natural burrow conditions at 28°–30°C. The animals were fed *ad libitum* daily on a diet of sweet potatoes, carrots, beetroot, apples and cucumber, and provided with wood wool (Vienna) or paper towel shreds (Pretoria) as nesting material. The boxes (toilet chamber) were cleaned once a day and the food container once a week. During observations, a standardised amount of digging substrate (1 x 200 ml wood shavings) was inserted into the tunnel system every 2 h to provide substrate for burrowing activity.

All individuals were identified via passive integrated transponder tags, and prior to observations, the individuals received unique colour marks applied with permanent markers. Sex was determined from the external genitalia (Pretoria) or via molecular sexing using buccal mucosa samples (Vienna). The breeding females were identified by their characteristic genital morphology. In Vienna, we were unable to identify the breeding males in the groups morphologically, and no sexual behaviour was observed during the study. Therefore, we included all individuals except for queens in the behavioural analysis as non-breeders.

### Data collection

Data from 169 non-breeding individuals (67 females, 102 males) were included in this study. In Vienna, data were collected from 72 animals between July 2018 and July 2019. Body mass was recorded a mean of 7.0 ± 1.5 times from every animal whenever the group was removed from the tunnel system (e.g. before observation sessions or when taking mucosa samples) by placing them on an electronic scale (accurate to the nearest gram). In Pretoria, data were collected from 97 animals in August 2020. Body mass measurements were taken once for each animal before their first observation. The mean body mass for all non-breeders was 38.3 ± 12.3 g (range 16-74 g), with 36.1 ± 11.6 g for females and 39.8 ± 12.5 g for males. Ages were known for 91 non-breeders in Pretoria and 8 non-breeders in Vienna. The mean age at the time of observation was 415.4 d, ranging from 140-1254 d.

Behavioural data were collected using instantaneous scan sampling. The behaviour of every animal in a group was recorded in 6-10-min intervals, depending on group size. In larger groups, 20 animals were arbitrarily chosen for observation, whereas in groups smaller than 20 individuals, all animals were included in the observation. The ethogram included 16 behaviours (Supplementary Table S1), and the observations were recorded on a handheld device using software Animal Behaviour Pro version 1.2 (University of Kent, UK). In Vienna, groups remained in their usual tunnel systems, whereas in Pretoria, they were transferred one day before observation to a tunnel system better suited for observations (Supplementary Figure S1). The animals were allowed 24 h to habituate to the observational tunnels.

Observation sessions lasted 6 h and were carried out between 08:00 and 16:00 by the same one or two observers that alternated every 30 min. The observational period was chosen because naked mole-rats show unpredictable activity patterns with considerable inter-individual variation (Riccio and Goldman 2000). In Vienna, each group was observed five times over a mean period of 216 ± 61 d, with a mean time of 54 ± 41 d between sessions. In Pretoria, each group was observed on three consecutive days. Over all 43 sessions, a mean of 161 ± 69 sampling events was recorded per individual (range 78-300, see Table 1).

**Table 1:**
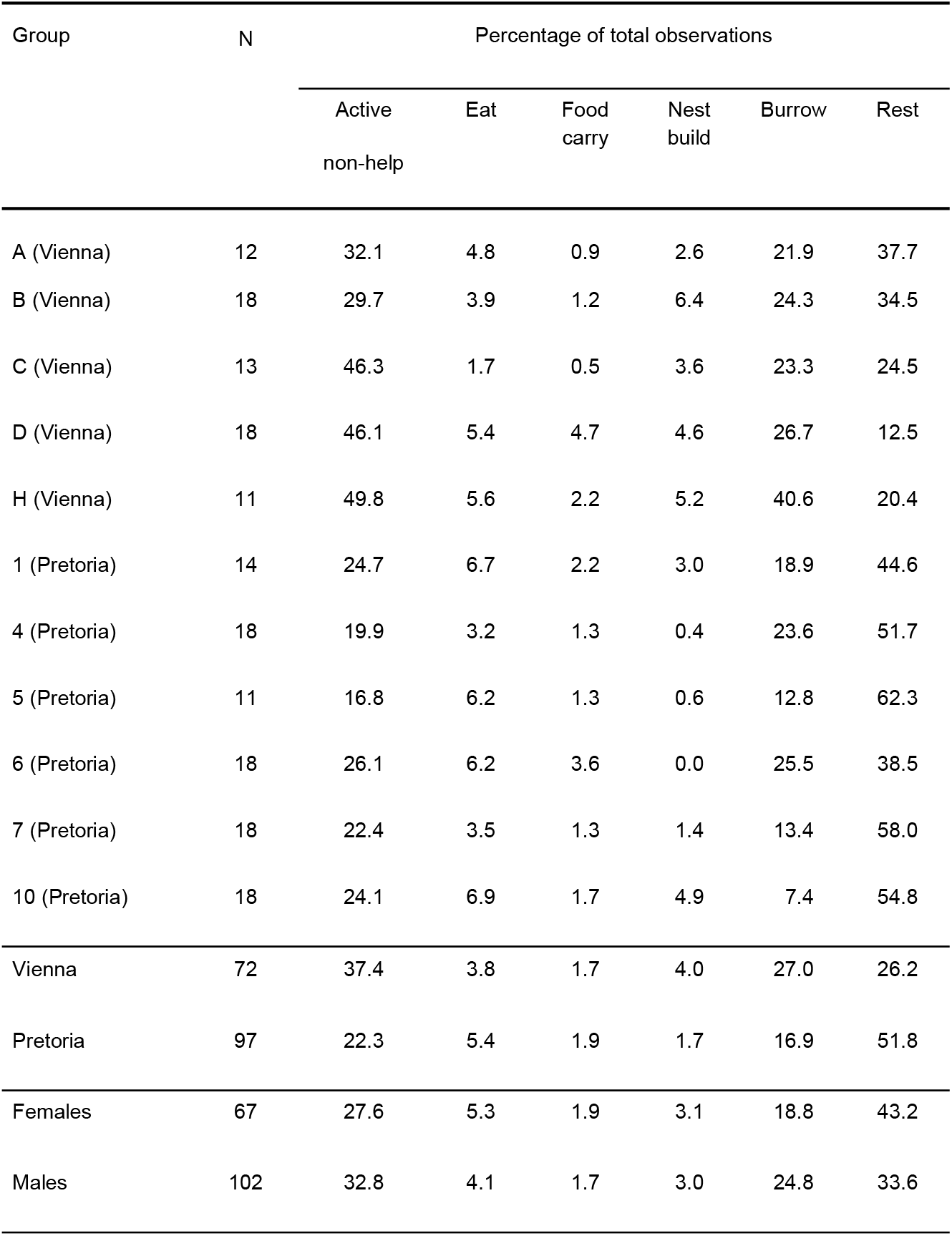
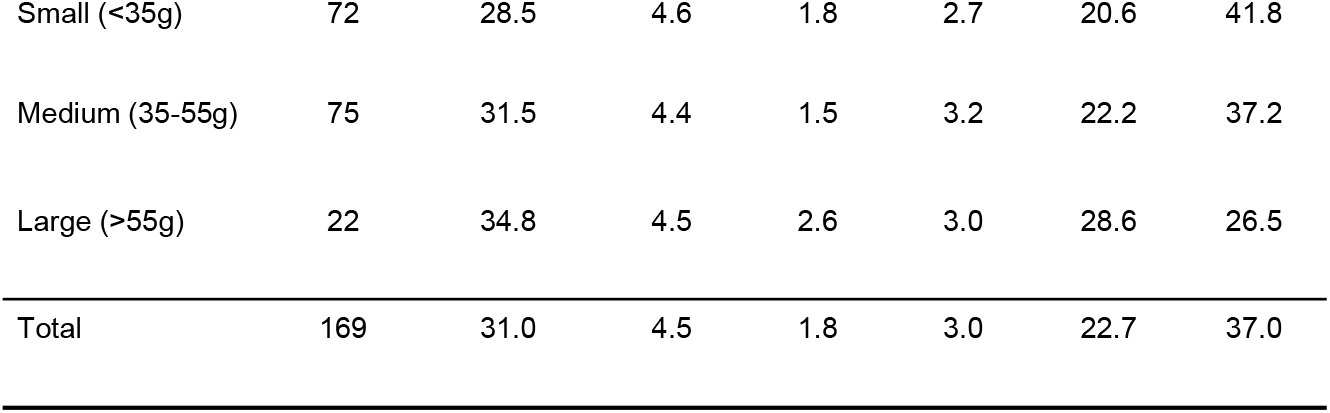
Descriptive summary statistics for the observational data used in this study split by group, population, sex and body mass. Population-level differences are also illustrated in Supplementary Figure S3.

### Statistical analysis

Individual correlations between types of cooperative behaviour and the effect of body mass and group size on cooperative behaviour were analysed with the use of three multilevel, multinomial behaviour models that increased in complexity due to the successive inclusion of fixed effect covariates and higher-level random effects (Koster and McElreath 2017). All three models were calculated separately for males and females. Subsequently, we also specified a model for a subset of animals of known age to investigate the effect of age on behaviour and a model for all non-breeders with sex as a predictor variable to quantify behavioural sex differences.

The 16 recorded behaviours were grouped into six categories: three types of cooperative behaviour, carrying food, nest building and burrowing (which aggregates all activities related to burrow maintenance such as gnawing at the tunnel walls, digging in or kicking and sweeping substrate), and three non-cooperative behaviour types, resting, eating and active non-help, which summarises all other active behaviours not related to cooperation so that distinction can be made between investment in cooperation and other activities. For most observations, no offspring were present, and the few recorded instances of pup carrying were excluded from the analyses.

The Widely Applicable Information Criterion (WAIC) was calculated to evaluate relative model fit, but due to their varying predictors and random effects structure, each of the three Models 1-3 and the comparison of their output provided information relevant to different aspects of our analysis of cooperative behaviour (Watanabe and Opper 2010, Watanabe 2013). The WAIC score was therefore not used for model selection, but rather as an indicator of model quality.

Model 1 included only intercepts and random effects for individuals and showed the extent of individual-level variance for each behavioural category as well as the within-individual correlations between the five non-resting behaviours. Since we were interested in individual trade-offs between active behaviours, resting was set as the reference category. This meant that coefficients of the intercepts indicated how much time individuals allocated to the respective behaviours relative to resting. Consequently, the variance of the reference category or correlations between the other behaviours and resting were not calculated.

In addition to the individual-level random effects, Model 2 included predictor variables that may be related to the expression of behavioural phenotypes in naked mole-rats. Body mass was added as a fixed effect to test the assumption that maximum body mass influences the cooperative investment of non-breeders. As another well-established predictor of behavioural contributions in cooperative societies the group size was also incorporated as a fixed covariate (Balshine et al. 2001, Fischer et al. 2014, Houslay et al. 2020). Both continuous predictors were z-score transformed before model fitting and specified as first- and second-order polynomials. To control for the origin of the population we also added this variable as a fixed factor with two levels (Vienna/Pretoria). The comparison of individual-level variances between Models 1 and 2 gave some indication of the proportion of variance in the behavioural categories that could be explained by the fixed effects. However, the inclusion of predictor variables can increase the higher-level variance estimates in multilevel models, which is why the variances in Model 2 should be interpreted with caution (Koster and McElreath 2017). The within-individual correlations between the behavioural responses are not sensitive to this issue, and the changes in correlation estimates relative to Model 1 reflected the impact of the predictor variables on the random effects.

The structure of Model 3 was further expanded to include random effects at the level of observation session and group, while maintaining the set of fixed effects from the previous model. Random effects for sessions were incorporated to account for temporal pseudo-replication created by recording the same individuals repeatedly throughout one session. Group-level random effects were introduced to adjust for clustering of the data by group. The complex random effects structure of this model affects the interpretation of the individual random effects and their correlations: individual-level variance estimates did not reflect variation across all the individuals of the population, but within-group variations and as a result, individual-level correlations in this model did not represent individual trade-offs between behavioural responses. However, including higher-level random effects improved the overall model fit and allowed a more precise estimation of the fixed effects. As a result, Model 3 was particularly suited for analysing the effects of the predictor variables on cooperative investment.

We expanded the structure of Model 3 to investigate the effect of age on behaviour for the subset of 99 non-breeders of known age by including age as a fixed effect (as a first-, second-and third-order polynomial) and litter as a random effect for Model 3a. Additionally, we applied Model 3b, which also retained the random and fixed effects structure of Model 3 but incorporated sex as a categorical predictor, to the whole dataset.

Models were fitted and analysed in a Bayesian framework with the R packages *rstan* (Stan Development Team 2020) and *rethinking* (McElreath 2020). Instead of the conventional Markov chain Monte Carlo algorithms, *rstan* employs Hamiltonian Monte Carlo chains, which are more efficient at achieving sufficiently mixed posterior distributions (Monnahan et al. 2017). We used three chains of 2000-3000 iterations for model fitting, half of which were devoted to the warm-up. To ensure adequate mixing of the chains, a non-centred parameterisation of the varying effects was realised with a Cholesky decomposition of the variance-covariance matrices (Koster and McElreath 2017). Additionally, we assigned weakly informative priors to the fixed effect parameters and variance-covariance matrices that prevent overfitting while influencing the posterior distribution as little as possible (Koster and McElreath 2017). To diagnose potential problems with chain mixing and convergence, we examined the trace plots and rank histograms of the chains as well as the effective number of samples and the Gelman-Rubin convergence diagnostic (R□ < 1.1) (McElreath 2020).

The correlations between random effects were considered significant if the 95% credible intervals of their posterior distributions did not include zero. The interpretation of the coefficients of the fixed effects is complicated because they do not represent the direct effect of the predictor on the probability of exhibiting a certain behaviour due to their relationship to the reference category. Following Koster and McElreath (2017), we instead calculated the predicted probabilities and their credible intervals in order to visualise the impact of body mass, group size and age on behaviour. Probabilities were based on fixed effects only while averaging over random effects. Prediction intervals cannot be used to test categorical predictor variables for significance, because they contain uncertainty from all covariates, and so to examine differences in behaviour between females and males, we calculated the contrasts between the predicted probabilities for the two groups (Koster and McElreath 2017). Statistical significance was inferred if the 95% credible intervals of the predicted differences did not span zero. All statistical analyses were performed in R (R Core Team, 2020).

### Ethical statement

The protocol used in this study was approved by the animal ethics committee of the University of Pretoria NAS099/2020 and Department of Agriculture land reform and rural development 12/11/1/8 (1595JD).

## RESULTS

### Individual-level trade-offs

Comparison with the WAIC showed that model fit improved with increased model complexity (Supplementary Table S2). Effects of the predictor variables are therefore presented for Model 3 and its variants 3a and 3b, while within-individual correlations are taken from Model 1 and 2.

We found no evidence of task specialisation of non-breeding naked mole-rats between any of the three cooperative tasks. Individual-level random effect correlations between any two of the observed behaviours (excluding the reference category resting) were positively correlated across both sexes and negative correlations were notably absent, indicating that there were no trade-offs between different cooperative behaviours within individuals (Table 2, Supplementary Tables S3 and S4 for random effects correlations on all levels from Models 1, 2 and 3 for females and males, respectively). Individuals that performed more of one cooperative behaviour were also more likely to engage in other cooperative behaviours: mole-rats who were more frequently observed carrying food engaged more often in nest building (females: ρ_3,4_ = 0.34 ± 0.13; males: ρ_3,4_ = 0.61 ± 0.08), and burrowing (females: ρ_3,5_ = 0.61 ± 0.10; males: ρ_3,5_ = 0.64 ± 0.07), while individuals who burrowed relatively more also allocated more of their time to nest building (females: ρ_4,5_ = 0.43 ± 0.11; males: ρ_4,5_ = 0.69 ± 0.06; values from Model 1, Table 2, upper half of each matrix). Most correlations remained robust after controlling for the influence of the fixed effects on behaviour in Model 2, though nest building was no longer significantly correlated to other cooperative tasks in females (Figure 1; Table 2, lower half of each matrix). The positive correlations remained qualitatively unchanged when limiting the dataset to include only individuals that were observed over a long period in the population from Vienna (Supplementary Figure S2, Supplementary Table S5). However, despite remaining positive, a small number of correlations, notably nest building to food carrying within the population in Vienna did not reach significance among females, presumably because the sample size was limited to 19 females.

**Table 2:** Correlations of individual-level random effects across responses from Model 1 and 2 for both sexes.

| Sex | Behaviour | Behaviour |  |  |  |  |
| --- | --- | --- | --- | --- | --- | --- |
|  |  | Active non-help | Eat | Food carry | Nest build | Burrow |
| Female | Active non-help | – | **0.59(0.09)** | **0.63(0.09)** | **0.58(0.09)** | **0.70(0.07)** |
|  | Eat | **0.56(0.13)** | – | **0.66(0.11)** | **0.61(0.11)** | **0.35(0.12)** |
|  | Food carry | **0.58(0.13)** | **0.58(0.14)** | – | **0.34(0.13)** | **0.61(0.10)** |
|  | Nest build | 0.22(0.14) | **0.45(0.15)** | 0.13(0.17) | – | **0.43(0.11)** |
|  | Burrow | **0.51(0.11)** | 0.24(0.14) | **0.51(0.13)** | 0.14(0.14) | – |
| Male | Active non-help | – | **0.60(0.08)** | **0.65(0.07)** | **0.78(0.05)** | **0.77(0.04)** |
|  | Eat | **0.64(0.08)** | – | **0.84(0.06)** | **0.57(0.09)** | **0.61(0.08)** |
|  | Food carry | **0.78(0.06)** | **0.83(0.06)** | – | **0.61(0.08)** | **0.64(0.07)** |
|  | Nest build | **0.66(0.07)** | **0.58(0.09)** | **0.70(0.08)** | – | **0.69(0.06)** |
|  | Burrow | **0.72(0.06)** | **0.60(0.08)** | **0.71(0.07)** | **0.58(0.09)** | – |

The upper half of the matrix lists correlations from Model 1, the lower half correlations from Model 2. Reported values are means from the posterior samples (SD in parenthesis); parameters in bold indicate estimates where the 95% credible intervals do not span zero.

**Figure 1:**
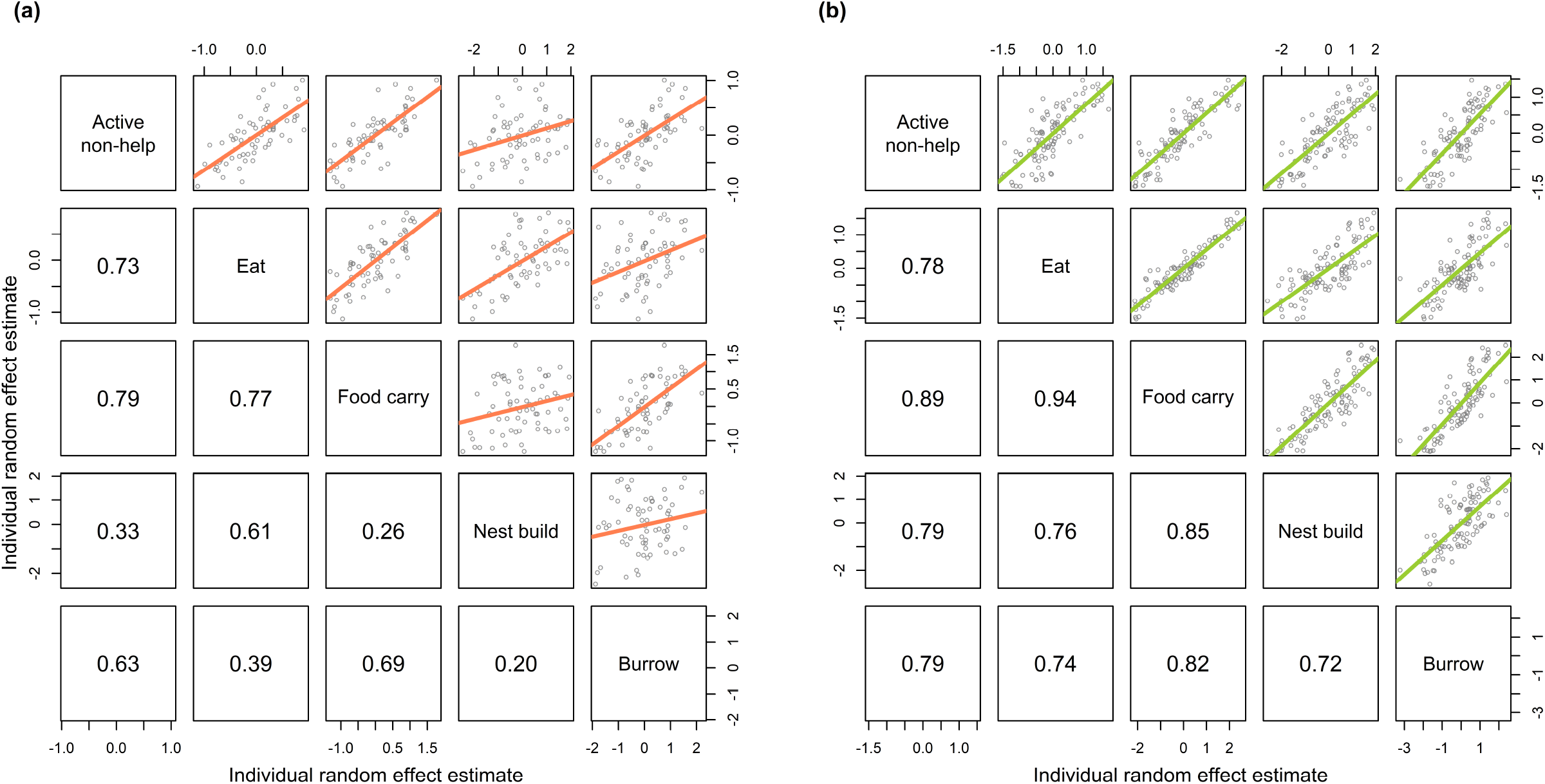
Within-individual random effects correlations from Model 2 for a) females and b) males. Values in the lower half of the matrix represent the correlations between the median individual level intercept in the posterior samples for each behaviour. They are therefore larger than the correlations presented in Table 2 that are taken directly from the variance-covariance matrices of the posterior samples.

### Effects of body mass, group size and age

Individual-level variances changed only to a small extent with the inclusion of fixed effects in Model 2 compared to Model 1, indicating that body mass, group size and population account for only a small proportion of individual-level behavioural variation (Supplementary Table S6). The behavioural changes attributed to the fixed effect estimates (body mass, group size, age) were also estimated with a high degree of uncertainty, suggesting that these individual and group characteristics were relatively poor predictors of cooperative behaviour of naked mole-rats in both populations. However, some attenuated general trajectories were notable in the visualisation of the predicted probabilities (Figures 2, 3 and 4).

**Figure 2:**
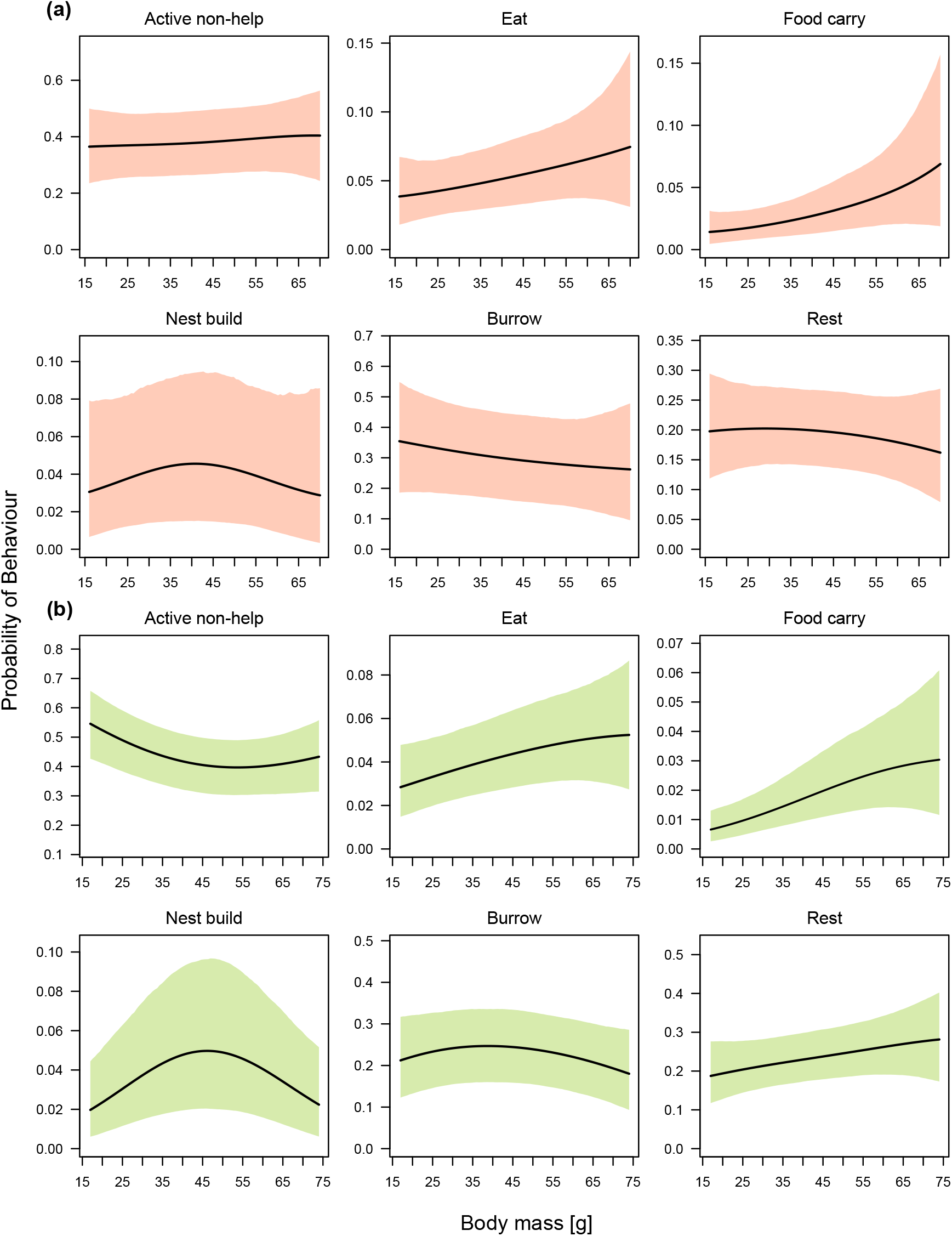
Model predictions of response behaviours as a function of body mass for **a)** females and **b)** males. All other fixed covariates are held at the sample mean and predictions are made at the population level for individuals from Vienna. Shaded regions show the 89% percentile intervals calculated from the posterior samples of Model 3 for each sex.

**Figure 3:**
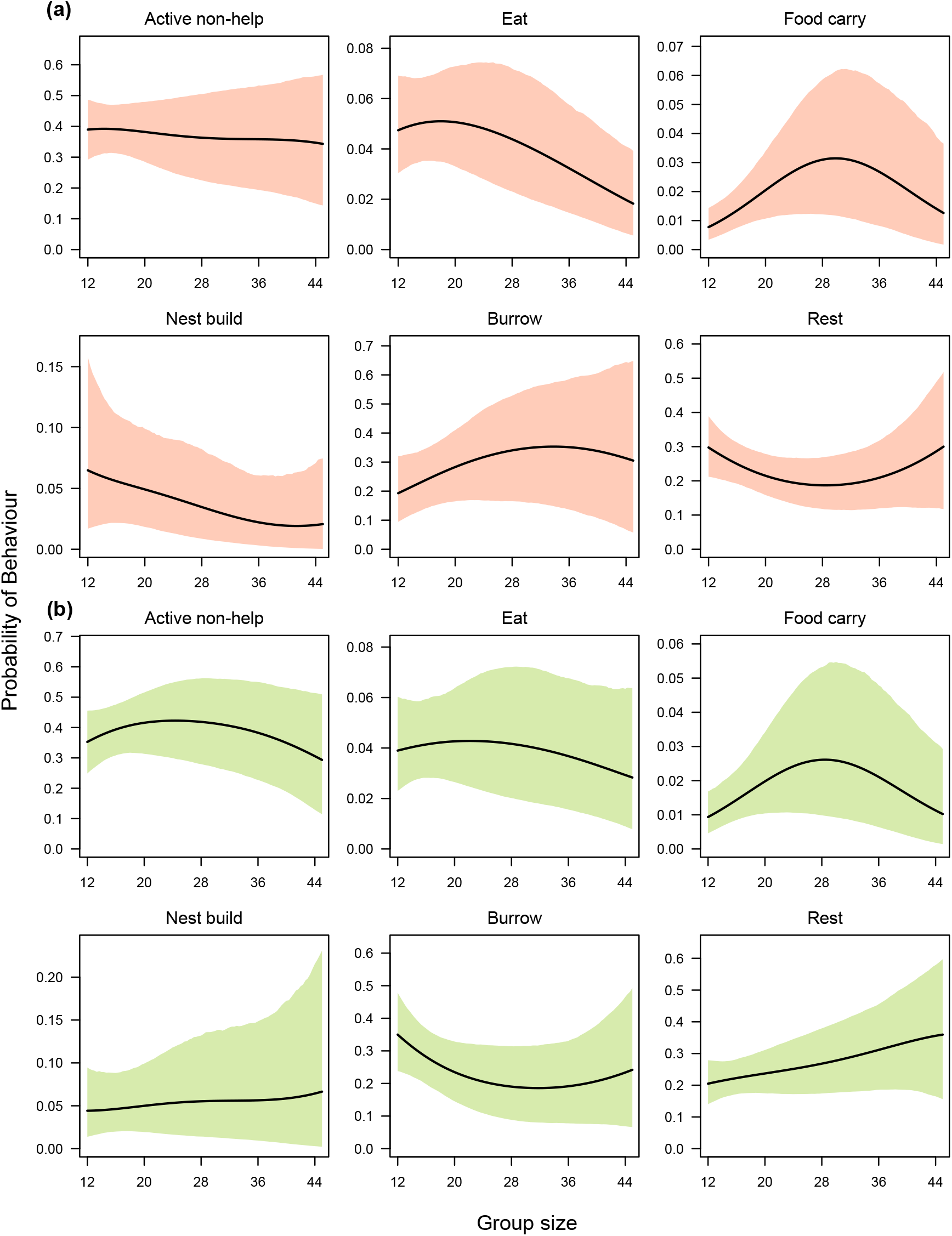
Model predictions of response behaviours as a function of group size for **a)** females and **b)** males. All other fixed covariates are held at the sample mean and predictions are made at the population level for individuals from Vienna. Shaded regions show the 89% percentile intervals calculated from the posterior samples of Model 3 for each sex.

**Figure 4:**
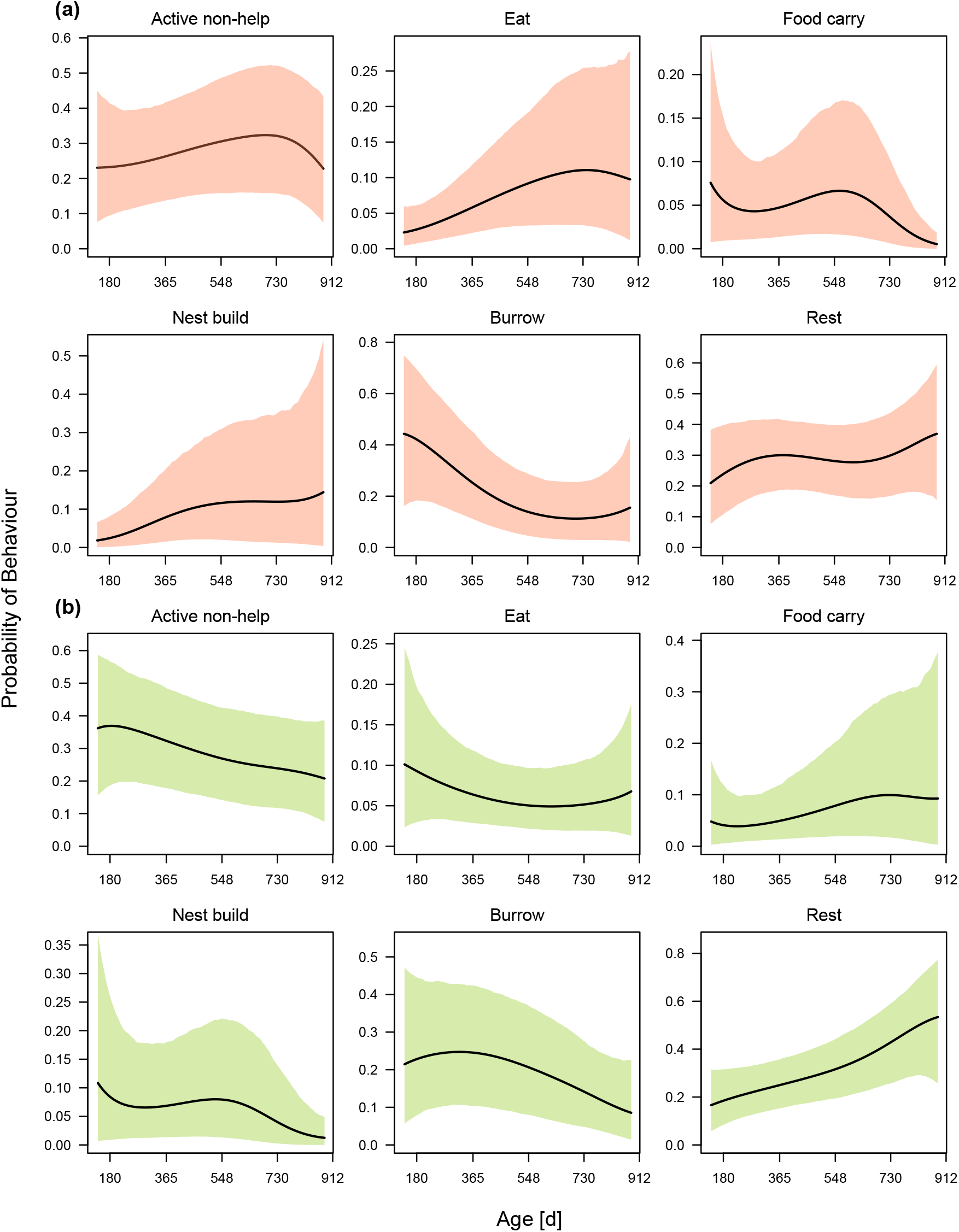
Model predictions of response behaviours as a function of age for **a)** females and **b)** males. All other fixed covariates are held at the sample mean. Shaded regions show the 89% percentile intervals calculated from the posterior samples of Model 3a for each sex.

Body mass had similar effects on male and female non-breeders (Figure 2, Supplementary Table S7). Food carrying increased with larger body mass, whereas burrowing activity decreased with larger body mass in both sexes. Individuals with intermediate body masses engaged mostly in nest-building activity. Group size had overall no convincing effect on the cooperative and non-cooperative behaviour of naked mole-rats (Figure 3, Supplementary Table S7). Individuals in larger groups rested more and furthermore showed less non-helping activity. Investment in food carrying peaked at intermediate group sizes in both sexes and burrowing showed a similar quadratic relationship in females. In contrast, burrowing declined and resting increased with larger group sizes in males. Females performed less nest building in larger groups, whereas no such trend was apparent for males.

Overall, the animals became less active with age, as seen by the increase of resting behaviour over time (Figure 4, Supplementary Table S7). In accordance with this trend, investment in burrowing behaviour decreased for both sexes during the first 2.5 years of life. The effect of age on some behaviours seems sex-dependent: time allocated to food carrying declined after 1.5 years in females, whereas in males, food carrying increased with age and reached its peak at around 2 years. Nest building behaviour showed the inverse trends, with males expressing nest-building more frequently with age and females displaying a steep decline after 1.5 years.

### Sex differences in cooperative behaviour

Whereas some sex differences existed in relation to age, overall, sex differences in cooperative and non-cooperative behaviour were minor and non-significant, with the exception of females spending marginally more time eating (Model 3b). Both sexes were equally likely to carry food, engage in nest building and burrowing after controlling for the effects of body mass, group size and population (Figure 5).

**Figure 5:**
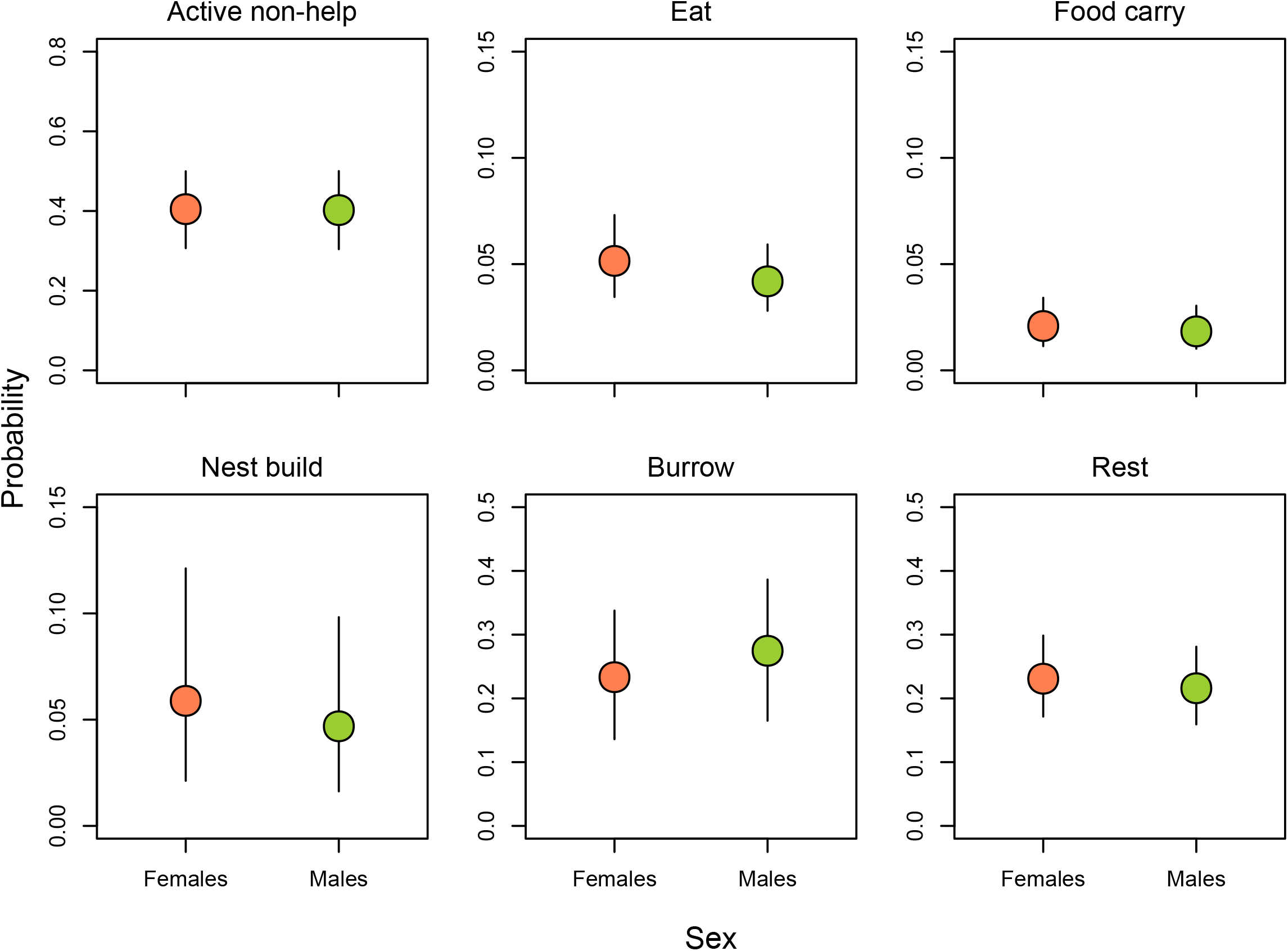
Model predictions of response behaviours as a function of sex (67 female non-breeders and 102 male non-breeders). All other fixed covariates are held constant at the sample mean. The confidence intervals are the 89% percentile intervals as calculated from the posterior samples of Model 3b.

## Discussion

Our results provide no indication that naked mole-rats in these captive groups specialise temporarily or permanently in specific work-related tasks and instead suggest that within individuals all work-related tasks correlate positively with each other. Overall, individuals show variation in total activity levels that trade-off against resting duration, whereas investment in all recorded behaviours, including burrowing, nest building and food carrying, correlate positively with each other. As such, naked mole-rat non-breeders vary in their general commitment to cooperative behaviour that may be linked to some differences in age, body size, metabolism and group size. Their behavioural types can be characterised along a one-dimensional syndrome varying from individuals that show long periods of activity and that engage frequently in all forms of work-related cooperative behaviour to individuals that show shorter periods of activity and engage less frequently in all forms of cooperative behaviour.

Our results do not support previous suggestions that naked mole-rats specialise on specific tasks, either temporarily or permanently (Jarvis 1981, Jarvis et al. 1991, Lacey and Sherman 1991, Lacey and Sherman 1997). These studies inferred specialisation from size-related variation in cooperative behaviour by showing that burrowing, nest building and food carrying follow divergent size-dependent trends. However, the individuals were mostly of unknown age and it has been found that size and age-related differences may not necessarily be the result of specialisation, but instead are often consequences of the relative costs and benefits to individuals at different life stages (McNamara and Houston 1996, Faulkes and Bennett 2016, Zöttl et al. 2016a, Gilbert et al. 2020). The results of our study also represent a clear example where size and age-related variation can exist despite a positive correlation of engagement in different tasks within individuals. One of the most important studies advancing the case that naked mole-rats may specialise on specific tasks used a principal component analysis to investigate whether individuals specialise across territory defence, pup care and work-related tasks and show that these three types of behaviours load on different axes (Mooney et al. 2015). However, specialisation at the individual level may be expected to result in the loading of different cooperative tasks with opposite directionality on the same axis and this has yet to be shown. Our study was unable to assess how territory defence and pup carrying relates to other cooperative tasks such as nest building, food carrying and burrowing because pup care behaviour is only shown in times when offspring are very small and territory defence needs to be elicited by introducing foreign conspecifics, predators or predator scent. Future research is now needed to clarify how these behaviours relate to each other in a similar analytical framework as we used in this study and to explicitly test whether the investment in pup care and territory defence are traded-off against investment in other tasks at the individual level.

Although the idea of specialization of non-breeders in mole-rat species is interesting and has attracted widespread attention, recent longitudinal studies suggest that individuals rarely specialise. Mole-rats within the genera *Fukomys* and *Cryptomys* show body size related changes in cooperative behaviour and individuals can vary widely in the frequency of burrowing behaviour (Bennett and Jarvis 1988, Bennett 1990, Burda 1990, Bennett 1992, Jarvis et al. 1994, Spinks et al. 1999, Scantlebury et al. 2006). However, longitudinal studies of Damaraland mole-rats of known ages have shown that individuals do not trade-off investment in cooperative behaviours and that the general patterns of the distribution of cooperative behaviour across individuals are similar to those of naked mole-rats shown in this study (Zöttl et al. 2016a, Thorley et al. 2018, Zöttl et al. 2018, Gilbert et al. 2020). Evidence from field studies of other *Fukomys* species also supports the notion that the behavioural similarity of mole-rats societies with obligatorily eusocial insects has probably been overemphasized in the past and evidence for specialisation, divergent developmental trajectories or bimodal trait distributions across individuals are rare (Faulkes and Bennett 2016, Šklíba et al. 2016, Zöttl et al 2016b, Van Daele et al. 2019, Voigt et al. 2019).

Our results are consistent with growing evidence suggesting that the distribution of cooperative behaviour across individuals in social mole-rat societies is similar to that of many other cooperatively breeding mammals and birds. In most cooperatively breeding mammals and birds, individuals vary in the overall commitment to cooperative behaviours. In meerkats and banded mongooses, for example, different helping behaviours are positively correlated to each other and divergent developmental trajectories are absent (Carter et al. 2014, Sanderson et al. 2015), though individuals vary by age, sex and opportunities to breed (Clutton-Brock et al. 2001, Clutton-Brock et al. 2003, Cant et al. 2016, Clutton-Brock and Manser 2016). Similarly, in cooperatively breeding birds, individuals show positively co-varying variation across individuals in specific cooperative tasks such as chick feeding and nest defence (van Asten et al. 2016, Teunissen et al. 2020) and although some notable exceptions exist (Arnold et al. 2005), the majority of studies suggest that across avian and mammalian cooperative breeders task specialisation is rare.

Whereas most research on cooperatively breeding mammals is often limited to observational data and relies on longitudinal studies of life-history variation, cooperatively breeding cichlids have emerged as some of the most promising and innovative study systems to investigate divergent developmental trajectories and individual specialisation among cooperatively breeding vertebrates (Arnold and Taborsky 2010, Bruintjes and Taborsky 2011, Taborsky et al. 2013, Fischer et al. 2017). The reproductive ecology of these fish shares many of the characteristics with cooperatively breeding birds and mammals. They live in stable groups with high reproductive skew and up to 25 helpers that engage in different forms of cooperative behaviour (Taborsky and Limberger 1981, Taborsky 1984, 1985). Helpers that vary in relatedness to the breeders can either stay in the territory and help or disperse and breed independently (Taborsky and Limberger 1981, Bergmuller et al. 2005, Dierkes et al. 2005, Heg et al. 2011, Hellmann et al. 2016), and cooperative behaviour is used to appease the dominant breeders and prevent punishment and eviction (Bergmuller and Taborsky 2005, Zöttl et al. 2013b, Fischer et al. 2014, Naef and Taborsky 2020). Experimental manipulations of the social environment and predation pressure have long-lasting effects on the behavioural phenotype and the physiology of helpers that may be related to variation life-histories and could adapt some individuals for extended philopatry and others to dispersal (Taborsky et al. 2013, Fischer et al. 2015, Fischer et al. 2017, Antunes et al. 2020), though it remains unclear whether divergent phenotypes are a result of adaptation, or of developmental constraints, and whether long-lasting developmental effects overshadow the capacity to adapt to stochastically arising breeding or dispersal opportunities (Bergmuller et al. 2005, Zöttl et al. 2013a, English et al. 2015). In contrast to predictions about the distribution of investment in specific tasks among specialised helpers, these cichlids show positive correlations between territory defence and maintenance tasks (Le Vin et al. 2011), and it remains unclear if developmental effects lead to some trade-offs between different tasks in cooperatively breeding cichlids.

Our study suggests that burrowing frequency in naked mole-rats decreased with age and body mass, which is consistent with recent research on age-related behavioural variation in naked mole-rats (Gilbert et al. 2020). Nest building peaked at intermediate body mass at the age of two years and food carrying increased in heavier individuals. These patterns are broadly similar to those in Damaraland mole-rats, and the general decline of burrowing behaviour coincides with the age and body mass at which individuals in the wild disperse from their natal group (Hochberg et al. 2016, Zöttl et al. 2016a, Torrents-Ticó et al. 2018a). While breeders and non-breeders show long lifespans in captivity (Dammann and Burda 2006, Buffenstein 2008, Schmidt et al. 2013, Ruby et al. 2018), most non-breeders in the wild disappear from their natal groups when they reach approximately 2-3 years of age, and subsequently either become breeders in a new group or die during dispersal (but see Young et al. 2015, Hochberg et al. 2016, Torrents-Ticó et al. 2018b). Those that remain in the group have been found to gain body mass and become less active (Jarvis et al. 1991, O’Riain et al. 1996, Thorley et al. 2018). This relationship is reflected in our groups, as male non-breeders rested more as they aged, and time spent on most cooperative activities peaked at around 1.5-2 years or generally decreased with age.

Our data reveal that sex differences in work-related cooperative behaviour of naked mole-rats are minor and body mass and age-related patterns are broadly similar in male and female non-breeding individuals. This is consistent with previous studies (Jarvis et al. 1991, Lacey and Sherman 1997, Gilbert et al. 2020), and it is possible that as in other social mole-rats species, sex-differences are limited to allo-parental care behaviour which we were unable to record in this study (Bennett 1990, Zöttl et al. 2016a, Thorley et al. 2018, Zöttl et al. 2018). The lack of sex-differences contrasts with the distribution of cooperative behaviours in many other cooperatively breeding species where sex differences are common and often linked to sex differences in philopatry (Clutton-Brock et al. 2002). In mole-rats the duration of philopatry differs only marginally between males and females (Braude 2000, Hazell et al. 2000, Torrents-Ticó et al. 2018b, Hochberg et al. 2016) and could be the underlying reason for similar investment in cooperative behaviour across both sexes in naked and other social mole-rat species. However, many sex differences in the behaviour of mammals only manifest after sexual maturity, and an alternative explanation for the lack of sex differences in naked mole-rat non-breeders is that non-breeders are hormonally pre-pubescent and therefore show little sex-specific variation (Faulkes et al. 1990, 1991 and 1994).

## Conclusion

Naked and Damaraland mole-rats have been proposed to share many of the traits of the obligatory eusocial insects, including task specialisation among workers. Our study suggests that task specialisation among different work-related tasks does not occur, which is consistent with recent studies on the distribution of cooperative behaviour among non-reproductive individuals of other mole-rat species. In contrast, individuals that contribute more to a specific task are also more likely to engage in a different task, suggesting that individuals primarily vary in their overall investment in cooperative behaviour. Our data suggest that naked mole-rats show similar behavioural organisation to other cooperatively breeding vertebrates where involvement in different tasks is commonly positively correlated within individuals and that similarity to the obligatorily eusocial insects has been overemphasized.

## Author contributions

MZ conceived the study. SS and RF collected the data with assistance from all authors. SS analysed the data. MZ and SS wrote the first draft of the paper. All authors commented and edited the manuscript.

## Significance statement

It has been controversially discussed whether non-breeders in naked mole-rats belong to distinct castes that specialise permanently or temporarily in specific cooperative tasks. In this paper we show that non-breeding individuals vary in overall cooperative investment, but do not specialise on specific work tasks. Our data provide no evidence for temporary specialisation and suggests that the behavioural organisation of naked mole-rats resembles that of other cooperatively breeding vertebrates more than that of eusocial insect species.

## Acknowledgments

We thank the Vienna Zoo for providing access to their mole-rat groups. We are particularly grateful to Anton Weissenbacher, Doris Preininger, Inez Walter and all caretakers for their help and support in this study. We thank Jack Thorley for helpful discussions about the analyses and interpretation of the data.

