## Supplemental materials for "Naked mole-rats (*Heterocephalus glaber*) do not specialise on cooperative tasks"

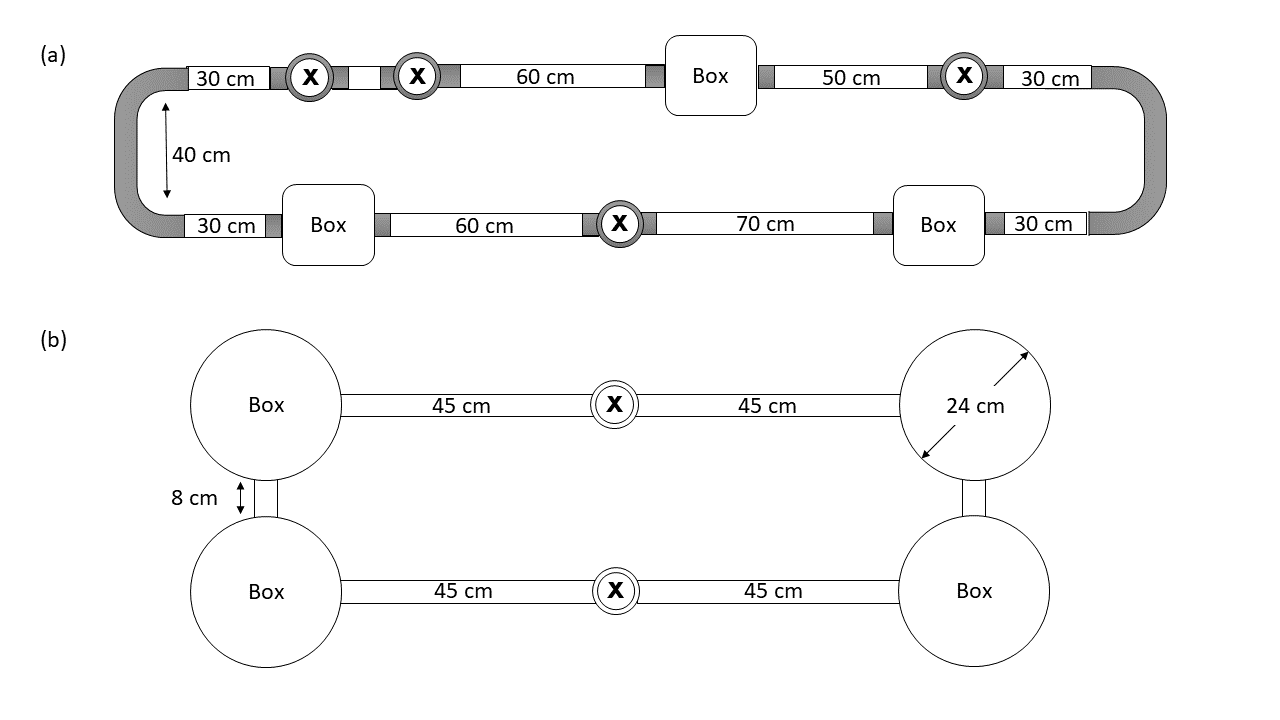

**Figure S1:** Schematic diagrams of a) a permanent tunnel system for 12 animals and b) the tunnel system used for observations at the University of Pretoria. Dimensions of the rectangle boxes are 15 x 15 x 15 cm and diameter of the tubes is 5 cm. Xs mark openings for the insertion of digging substrate.

**Table S1**: Naked mole-rat ethogram

| Response category | | Subcategories | Description |
| --- | --- | --- | --- |
| Active  non-help | | Locomotion | Moving through the tunnel system, not engaged in obvious work. |
|  |  | Other | Behaviours that cannot be assigned to other categories or the subject but not its behaviour can be identified. |
|  |  | Self-groom | Self-directed grooming, e.g. scratching, wiping, cleaning own body with incisors and feet. |
|  |  | Sniff | Sniffing objects, the tubes, the air or other individuals. |
|  |  | Social interaction | “Incisor fencing” (locking incisors, shoving each other back and forth), biting, nuzzling each other’s’ bodies and genitals. |
| Eat | | Eat | Eating food. |
| Food carry | (cooperation) | Food carry | Transporting food by pushing or dragging it through the tubes. |
| Nest build | (cooperation) | Nest build | Engaging with nest material (wood wool) by either dragging it in the direction of the nest or trying to pull it out of a certain location, but not sweeping it. |
| Pup carry | (cooperation) | Pup carry | Carrying or dragging pups through the tunnels with the incisors. |
| Burrow | (cooperation) | Dig | Using incisors and forelegs to dig in the litter or attempting to dig at the tube. |
|  |  | Gnaw | Scratching, biting, chewing on the tunnel walls with the incisors. |
|  |  | Volcano | Kicking litter upwards out of the tubes or boxes with the hindlegs. In contrast to sweeping, material is moved vertically, and the animal does not move backward between successive kicks. |
|  |  | Locomotion between sweep | Moving between bouts of sweeping. |
|  |  | Sweep | Moving backwards while pushing litter with the hind legs. |
| Rest | | Rest | Subject is immobile in a tunnel, head is down. |
|  |  | Sleep | Subject lies in the nest box with its eyes closed. |

**Table S2:** Comparison of Models 1, 2 and 3 for both sexes with WAIC (widely applicable information criterion)

| Sex | Model | Random effects | Fixed effects | WAIC (SE) | ΔWAIC(SE) | Weight |
| --- | --- | --- | --- | --- | --- | --- |
| Female |  |  |  |  |  |  |
|  | 1 | Individual | - | 23342.7(158.36) | 689.9(52.37) | 0 |
|  | 2 | Individual | Group size^1^,  body mass^1^,  population | 23334.4(159.23) | 681.5(51.73) | 0 |
|  | 3 | Individual,  scan, group | Group size^1^,  body mass^1^, population | 22652.9(163.92) | - | 1 |
| Male |  |  |  |  |  |  |
|  | 1 | Individual | - | 44816.2(211.44) | 1487.2(74.57) | 0 |
|  | 2 | Individual | Group size^1^, body mass^1^,  population | 44760.6(212.32) | 1431.6(72.94) | 0 |
|  | 3 | Individual,  scan, group | Group size^1^, body mass^1^, population | 43329.0(218.53) | - | 1 |

1 Included as first- and second-order polynomials

**Table S3**: Correlations (SD in parentheses) of random effects across the behavioural responses in Models 1-3 for females

| Sex | Model, | Behaviour | Behaviour | | | | | | | | | |
| --- | --- | --- | --- | --- | --- | --- | --- | --- | --- | --- | --- | --- |
|  | random effect |  | Active  non-help | | Eat | | Food  carry | | Nest build | | Burrow | |
| Female | 1, | Active non-help | |  | | **0.59(0.09)** | | **0.63(0.09)** | | **0.58(0.09)** | | **0.70(0.07)** |
|  | Individual | Eat | | - | |  | | **0.66(0.11)** | | **0.61(0.11)** | | **0.35(0.12)** |
|  | level | Food carry | | - | | **-** | |  | | **0.34(0.13)** | | **0.61(0.10)** |
|  |  | Nest build | | - | | - | | - | |  | | **0.43(0.11)** |
|  |  | Burrow | | - | | - | | - | | - | |  |
| Female | 2, individual level | Active non-help | |  | | **0.56(0.13)** | | **0.58(0.13)** | | 0.22(0.14) | | **0.51(0.11)** |
|  |  | Eat | | - | |  | | **0.58(0.14)** | | **0.45(0.15)** | | 0.24(0.14) |
|  |  | Food carry | | - | | - | |  | | 0.13(0.17) | | **0.51(0.13)** |
|  |  | Nest build | | - | | - | | - | |  | | 0.14(0.14) |
|  |  | Burrow | | - | | **-** | | **-** | | **-** | |  |
| Female | 3, individual level | Active non-help | |  | | **0.50(0.14)** | | **0.51(0.16)** | | 0.34(0.17) | | **0.58(0.11)** |
|  |  | Eat | | - | |  | | **0.50(0.17)** | | 0.39(0.19) | | 0.25(0.15) |
|  |  | Food carry | | - | | - | |  | | 0.25(0.20) | | **0.50(0.15)** |
|  |  | Nest build | | - | | - | | - | |  | | **0.46(0.15)** |
|  |  | Burrow | | - | | - | | - | | - | |  |
| Female | 3,  scan  level | Active non-help | |  | | **0.63(0.13)** | | **0.54(0.15)** | | **0.61(0.14)** | | **0.71(0.10)** |
|  |  | Eat | | - | |  | | **0.76(0.12)** | | **0.65(0.15)** | | **0.64(0.14)** |
|  |  | Food carry | | - | | - | |  | | **0.52(0.17)** | | **0.54(0.15)** |
|  |  | Nest build | | - | | - | | - | |  | | **0.70(0.12)** |
|  |  | Burrow | | - | | - | | - | | - | |  |
| Female | 3,  group level | Active non-help | |  | | 0.02(0.36) | | 0.03(0.35) | | -0.03(0.34) | | 0.02(0.35) |
|  |  | Eat | | - | |  | | 0.06(0.36) | | 0.08(0.34) | | 0.01(0.35) |
|  |  | Food carry | | - | | - | |  | | -0.01(0.35) | | 0.03(0.35) |
|  |  | Nest build | | - | | - | | - | |  | | -0.04(0.29) |
|  |  | Burrow | | - | | - | | - | | - | |  |

Estimates represent the means from the posterior samples (SD in parentheses). Parameters in bold indicate estimates where the 95% credible intervals do not span zero.

**Table S4**: Correlations (SD in parentheses) of random effects across the behavioural responses in Models 1-3 for males

| Sex | Model, | Behaviour | | Behaviour | | | | | | | | |
| --- | --- | --- | --- | --- | --- | --- | --- | --- | --- | --- | --- | --- |
|  | random effect |  | | Active  non-help | | Eat | | Food carry | | Nest build | | Burrow |
| Male | 1, | Active non-help |  | | **0.60(0.08)** | | **0.65(0.07)** | | **0.78(0.05)** | | **0.77(0.04)** | |
|  | Individual | Eat | - | |  | | **0.84(0.06)** | | **0.57(0.09)** | | **0.61(0.08)** | |
|  | level | Food carry | - | | - | |  | | **0.61(0.08)** | | **0.64(0.07)** | |
|  |  | Nest build | - | | - | | - | |  | | **0.69(0.06)** | |
|  |  | Burrow | - | | - | | - | | - | |  | |
| Male | 2, individual level | Active non-help |  | | **0.64(0.08)** | | **0.78(0.06)** | | **0.66(0.07)** | | **0.72(0.06)** | |
|  |  | Eat | - | |  | | **0.83(0.06)** | | **0.58(0.09)** | | **0.60(0.08)** | |
|  |  | Food carry | - | | - | |  | | **0.70(0.08)** | | **0.71(0.07)** | |
|  |  | Nest build | - | | - | | - | |  | | **0.58(0.09)** | |
|  |  | Burrow | - | | - | | - | | - | |  | |
| Male | 3, individual level | Active non-help |  | | **0.73(0.08)** | | **0.79(0.08)** | | **0.68(0.08)** | | **0.71(0.06)** | |
|  |  | Eat | - | |  | | **0.74(0.10)** | | **0.66(0.10)** | | **0.66(0.09)** | |
|  |  | Food carry | - | | - | |  | | **0.75(0.09)** | | **0.77(0.07)** | |
|  |  | Nest build | - | | - | | - | |  | | **0.81(0.06)** | |
|  |  | Burrow | - | | - | | - | | - | |  | |
| Male | 3,  scan  level | Active non-help |  | | **0.66(0.12)** | | **0.65(0.12)** | | **0.67(0.11)** | | **0.79(0.07)** | |
|  |  | Eat | - | |  | | **0.72(0.11)** | | **0.64(0.13)** | | **0.55(0.13)** | |
|  |  | Food carry | - | | - | |  | | **0.68(0.13)** | | **0.64(0.13)** | |
|  |  | Nest build | - | | - | | - | |  | | **0.60(0.13)** | |
|  |  | Burrow | - | | - | | - | | - | |  | |
| Male | 3,  group level | Active non-help |  | | -0.06(0.34) | | 0.06(0.34) | | 0.14(0.31) | | 0.24(0.32) | |
|  |  | Eat | - | |  | | 0.26(0.36) | | -0.04(0.32) | | 0.01(0.33) | |
|  |  | Food carry | - | | - | |  | | 0.04(0.33) | | 0.06(0.33) | |
|  |  | Nest build | - | | - | | - | |  | | -0.27(0.29) | |
|  |  | Burrow | - | | - | | - | | - | |  | |

Estimates represent the means from the posterior samples (SD in parentheses). Parameters in bold indicate estimates where the 95% credible intervals do not span zero.

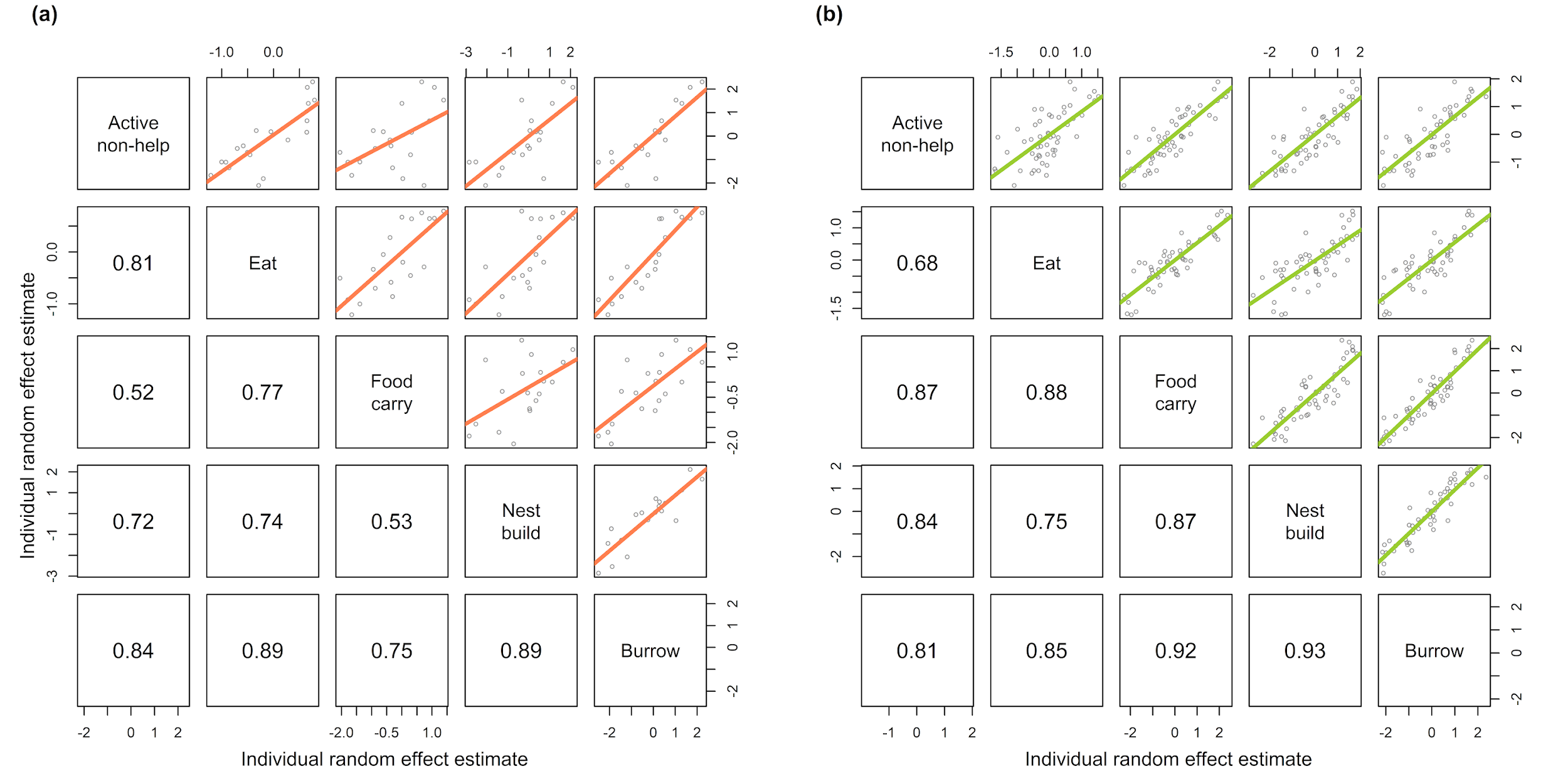
**Fig. S2**: Within-individual random effects correlations from Model 2 for **a)** females (n = 19) and **b)** males (n = 48) from the five groups observed in Vienna. Values in the lower half of the matrix represent the correlations between the median individual level intercept in the posterior samples for each behaviour. They are therefore larger than the correlations presented in Tab. S5 that are taken directly from the variance-covariance matrices of the posterior samples.

**Table S5.** Correlations (SD in parenthesis) of individual-level random effects across responses from Model 1 and 2 for both females and males from Vienna only

| Sex | Behaviour | Behaviour | | | | |
| --- | --- | --- | --- | --- | --- | --- |
|  |  | Active  non-help | Eat | Food carry | Nest build | Burrow |
| Female | Active non-help | − | **0.47(0.20)** | **0.39(0.20)** | 0.12(0.20) | **0.41(0.17)** |
|  | Eat | **0.48(0.23)** | − | 0.52(0.21) | 0.30(0.21) | **0.49(0.19)** |
|  | Food carry | 0.21(0.25) | 0.42(0.23) | − | 0.26(0.21) | 0.36(0.20) |
|  | Nest build | 0.42(0.21) | 0.37(0.24) | 0.20(0.25) | − | **0.60(0.16)** |
|  | Burrow | **0.62(0.15)** | 0.20(0.25) | **0.47(0.21)** | **0.67(0.15)** | − |
| Male | Active non-help | − | **0.49(0.12)** | **0.65(0.10)** | **0.72(0.08)** | **0.70(0.07)** |
|  | Eat | **0.47(0.21)** | − | **0.75(0.09)** | **0.52(0.12)** | **0.58(0.11)** |
|  | Food carry | **0.72(0.10)** | **0.69(0.12)** | − | **0.60(0.11)** | **0.63(0.10)** |
|  | Nest build | **0.71(0.09)** | **0.53(0.13)** | **0.69(0.11)** | − | **0.75(0.07)** |
|  | Burrow | **0.70(0.08)** | **0.69(0.09)** | **0.78(0.08)** | **0.82(0.06)** | − |

The upper half of the matrix lists correlations from Model 1, the lower half correlations from Model 2. Reported values are means from the posterior samples (SD in parenthesis); parameters in bold indicate estimates where the 95% credible intervals do not span zero.

**Table S6:** Variance estimates (SD in parenthesis) of the random effects for Model 1, 2 and 3 of each sex

| Random effect |  | Female | | | | | |  | | Male | | | | |
| --- | --- | --- | --- | --- | --- | --- | --- | --- | --- | --- | --- | --- | --- | --- |
|  |  | Model 1 | | Model 2 | | Model 3 | |  | | Model 1 | | Model 2 | | Model 3 |
| Individual level |  |  |  | |  | |  | |  | |  | |  | |
| Active non-help |  | 0.74(0.07) | 0.48(0.06) | | 0.44(0.05) | |  | | 0.94(0.06) | | 0.75(0.06) | | 0.57(0.05) | |
| Eat |  | 0.73(0.09) | 0.59(0.08) | | 0.52(0.08) | |  | | 0.78(0.07) | | 0.76(0.07) | | 0.60(0.07) | |
| Food carry |  | 1.17(0.14) | 0.90(0.14) | | 0.77(0.15) | |  | | 1.30(0.12) | | 1.23(0.11) | | 0.94(0.11) | |
| Nest build |  | 1.81(0.22) | 1.40(0.23) | | 0.85(0.16) | |  | | 1.37(0.12) | | 1.18(0.11) | | 1.04(0.10) | |
| Burrow |  | 1.29(0.12) | 1.00(0.11) | | 0.84(0.09) | |  | | 1.14(0.08) | | 1.08(0.08) | | 0.88(0.07) | |
| Scan level |  |  |  | |  | |  | |  | |  | |  | |
| Active non-help |  |  |  | | 0.69(0.08) | |  | |  | |  | | 0.70(0.08) | |
| Eat |  |  |  | | 0.60(0.09) | |  | |  | |  | | 0.65(0.09) | |
| Food carry |  |  |  | | 1.08(0.17) | |  | |  | |  | | 1.02(0.15) | |
| Nest build |  |  |  | | 0.90(0.15) | |  | |  | |  | | 0.82(0.11) | |
| Burrow |  |  |  | | 0.61(0.08) | |  | |  | |  | | 0.62(0.08) | |
| Group level |  |  |  | |  | |  | |  | |  | |  | |
| Active non-help |  |  |  | | 0.16(0.14) | |  | |  | |  | | 0.40(0.21) | |
| Eat |  |  |  | | 0.17(0.14) | |  | |  | |  | | 0.32(0.21) | |
| Food carry |  |  |  | | 0.29(0.25) | |  | |  | |  | | 0.47(0.32) | |
| Nest build |  |  |  | | 1.40(0.51) | |  | |  | |  | | 1.21(0.49) | |
| Burrow |  |  |  | | 0.73(0.32) | |  | |  | |  | | 0.55(0.24) | |

The reported quantities are the standard deviations of the random effects while the values in parentheses are the standard deviations of these quantities in the posterior samples.

**Table S7**: Posterior means (SD in parentheses) of fixed effects in Models 3 and 3a for each sex

| Model | Fixed effect | Active  non-help | Eat | Food carry | Nest build | Burrow |
| --- | --- | --- | --- | --- | --- | --- |
| 3, females | Group size | **−**0.16(0.22) | 0.07(0.23) | 0.82(0.37) | −0.25(0.52) | 0.49(0.37) |
|  | Group size^2^ | **−**0.16(0.13) | −0.31(0.14) | **−0.62(0.23)** | −0.33(0.37) | −0.31(0.24) |
|  | Body mass | 0.04(0.07) | 0.17(0.10) | 0.36(0.16) | 0.09(0.17) | −0.05(0.12) |
|  | Body mass^2^ | 0.02(0.05) | 0.02(0.07) | 0.01(0.10) | −0.09(0.13) | 0.02(0.08) |
|  | Population | **−1.27(0.31)** | **−**0.44(0.31) | **−**1.12(0.49) | **−**1.54(0.71) | **−1.60(0.58)** |
| 3a, females | Age | **−**0.15(0.25) | 0.00(0.23) | 0.72(0.39) | −0.29(0.56) | 0.60(0.38) |
|  | Age^2^ | **−**0.16(0.14) | −0.28(0.14) | **−0.56(0.23)** | −0.27(0.38) | −0.35(0.23) |
|  | Age^3^ | 0.04(0.07) | 0.16(0.10) | 0.36(0.16) | 0.09(0.18) | −0.05(0.12) |
| 3, males | Group size | −0.03(0.30) | −0.10(0.31) | 0.44(0.42) | −0.03(0.55) | −0.51(0.38) |
|  | Group size^2^ | **−**0.07(0.12) | **−**0.07(0.12) | **−**0.28(0.17) | **−**0.07(0.24) | 0.11(0.15) |
|  | Body mass | **−0.17(0.08)** | 0.06(0.09) | **0.28(0.14)** | 0.02(0.14) | **−**0.11(0.10) |
|  | Body mass^2^ | 0.05(0.05) | **−**0.02(0.06) | **−**0.07(0.09) | −0.17(0.09) | **−**0.04(0.06) |
|  | Population | **−1.11(0.39)** | −0.20(0.36) | −0.28(0.51) | **−1.63(0.68)** | −0.77(0.46) |
| 3a, males | Age | −0.08(0.33) | −0.16(0.29) | 0.31(0.42) | −0.05(0.58) | −0.58(0.38) |
|  | Age^2^ | **−**0.07(0.13) | **−**0.05(0.12) | **−**0.25(0.17) | **−**0.05(0.25) | 0.12(0.15) |
|  | Age^3^ | **−0.16(0.08)** | 0.06(0.09) | **0.29(0.14)** | 0.02(0.14) | **−**0.11(0.11) |

Parameters in bold indicate estimates whose 95% credible intervals do not span zero.

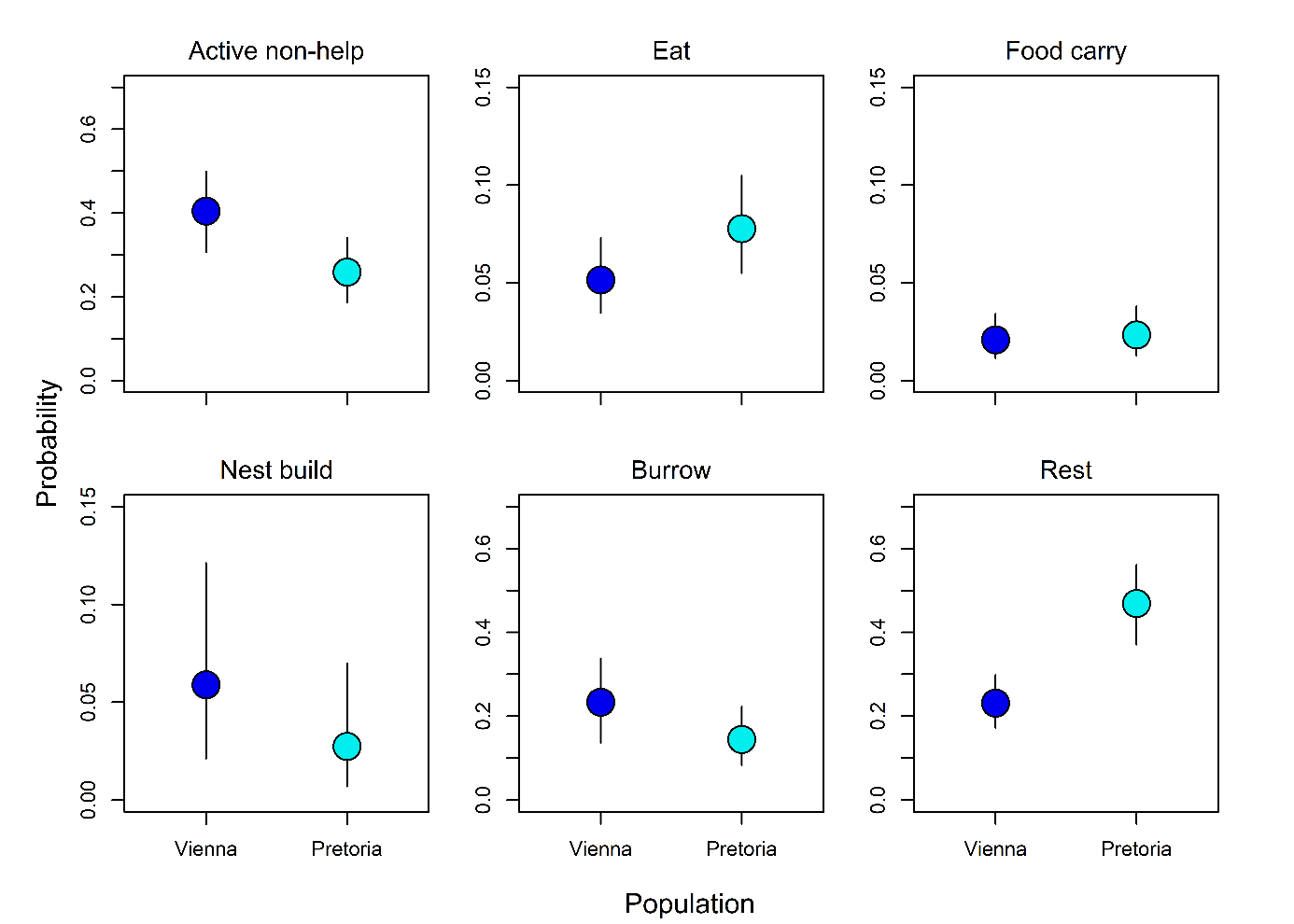

**Figure S3:** Model predictions of response behaviours as a function of population (72 non-breeders from Vienna and 97 non-breeders from Pretoria). All other fixed covariates are held constant at the sample mean. The confidence intervals are the 89% percentile intervals as calculated from the posterior samples of Model 3b. Credible intervals of the predicted differences showed that only resting behaviour differed significantly between the two populations.
